# BDNF stimulates the retrograde pathway for axonal autophagy

**DOI:** 10.1101/2022.08.08.503181

**Authors:** David Sidibe, Vineet Vinay Kulkarni, Audrey Dong, Jessica Brandt Herr, Sandra Maday

**Author notes:** Co-first authors (David Sidibe, Vineet Vinay Kulkarni).

## Abstract

Autophagy is a lysosomal degradative pathway important for neuronal development, function, and survival. But how autophagy in axons is regulated by neurotrophins to impact neuronal viability and function is poorly understood. Here, we investigate the regulation of axonal autophagy by the neurotrophin BDNF, and elucidate whether autophagosomes carry BDNF-mediated signaling information. We find that BDNF induces autophagic flux in primary neurons by stimulating the retrograde pathway for autophagy in axons. We observed an increase in autophagosome density and retrograde flux in axons, and a corresponding increase in autophagosome density in the soma. However, we find little evidence of autophagosomes co-migrating with BDNF. In contrast, BDNF effectively engages its cognate receptor TrkB to undergo retrograde transport in the axon. These compartments, however, are distinct from LC3-positive autophagic organelles in the axon. Together, we find that BDNF stimulates autophagy in the axon, but retrograde autophagosomes do not appear to carry neurotrophin-based signaling information. Thus, BDNF likely stimulates autophagy as a consequence of BDNF-induced processes that require canonical roles for autophagy in degradation, rather than unconventional roles in neurotrophin signaling.

## INTRODUCTION

Autophagy is a lysosomal degradation pathway that regulates neuronal development, function, and survival, which has important consequences for the axonal compartment. In fact, autophagy regulates axonal outgrowth and guidance (1-4), and loss of key autophagy genes can alter the formation of axon tracts connecting left and right hemispheres in the murine brain (2,3,5). Some reports indicate that loss of bulk autophagy increases axon length (1,4), suggesting that autophagy restrains axonal outgrowth. By contrast, loss of selective autophagy, and not bulk autophagy, does not alter axon growth or branching, but rather axon guidance (2). These effects are mediated through a combination of mechanisms that reduce axonal responsiveness to guidance cues, and non-cell autonomous effects of mislocalized glial guideposts, which result in reduced interhemispheric tracts and increased axonal mistargeting *in vivo* (2). Moreover, autophagy facilitates the assembly of presynaptic compartments during synaptogenesis (4,6). Clearly, autophagy serves key roles in establishing proper connectivity of the nervous system.

Autophagy is also important in maintaining axonal homeostasis and function. Several reports find that autophagy negatively regulates presynaptic neurotransmission by reducing synaptic vesicle pool size and release probability (7-11). Loss of autophagy also results in degeneration of established neuronal networks and neuromuscular junctions (5,12-18). Neuron-specific deletion of core autophagy genes is sufficient to cause swelling of the distal axon followed by retraction and decay (14-16), indicating cell autonomous roles for autophagy in maintaining axonal homeostasis. The extent to which developmental defects may contribute to the degeneration observed postnatally in these mouse models is unclear. Combined, the varied phenotypes elicited by the loss of autophagy depend on the type of autophagy examined, developmental stage, neuronal compartment, neuronal subtype, and contributions from neighboring glia. Thus, it is of critical interest to define the mechanisms and regulation of autophagy in neuronal axons that enable these diverse functions.

During autophagy, components of the cytoplasm (e.g. proteins and organelles) are encapsulated within a membrane that fuses with itself to form a double membraned autophagosome. Autophagosomes then undergo a series of fusion events with endolysosomal organelles to mature into autolysosomes that enable cargo degradation (19-21). Detection of double-membraned autophagosomes in axons dates back to classic electron microscopy studies (22,23). Subsequent live cell imaging studies revealed the dynamic nature of autophagy in axons in real time. Autophagosomes in axons are largely produced in the distal axon and presynaptic terminals (4,23-28). Following formation, autophagosomes undergo retrograde transport driven by dynein that delivers these organelles to the soma where they fuse with lysosomes to fully mature into degradative autolysosomes (23,24,29-31). The journey to the soma involves a series of maturation steps in the axon involving fusion with LAMP1-positive organelles, to yield amphisomes, that deliver lysosomal components and degradative enzymes to axonal autophagosomes, and may confer some degree of proteolytic activity (24,29,31-33). In sum, axonal autophagy is a vectorial pathway that delivers cargo from the distal axon to the soma, which may facilitate encounters with degradative lysosomes enriched in this region of the neuron. Importantly, this pathway is conserved in intact nervous systems in vivo (27,34-36), indicating a foundational pathway to maintain axonal homeostasis. But how autophagy in axons is regulated to impact neuronal viability and function is poorly understood and remains a major area of investigation.

One source of regulation may come from neurotrophins. Neurotrophins are a family of growth factors secreted by target tissue that regulate the development, function, and survival of innervating neurons (37,38). Neurotrophins bind to receptors on the presynaptic membrane (e.g. Brain-Derived Growth Factor [BDNF] binds to its receptor tropomyosin-related kinase B [TrkB]), and are endocytosed to form signaling endosomes. These signaling endosomes then undergo retrograde transport in the axon to reach the soma and elicit changes in gene expression (37-41). Several lines of evidence suggest that BDNF regulates autophagy in neurons (42,43). Kononenko et al. report that treatment of primary cortico-hippocampal neurons with BDNF increases the speed of retrograde autophagosomes in the axon, but effects on autophagy levels are unknown (43). In fact, since autophagosomes and signaling endosomes both undergo retrograde transport (24,39), several models have proposed that these organelle populations may be merging. Indeed, several groups have shown colocalization between axonal autophagosomes and TrkB (43,44). However, it is unclear what proportion of autophagosomes contain TrkB. Nonetheless, these data raise the possibility that retrograde axonal autophagosomes may carry signaling information in the form of BDNF that may be important for neuronal development and function (43,44). By contrast, Wang et al. show that blocking activation of TrkB in primary neurons does not affect the number of retrograde autophagosomes in the axon, suggesting that TrkB signaling is not required for constitutive autophagic flux in axons (45). Moreover, other studies find an inhibitory effect of BDNF on autophagic flux (42,46). In fact, Nikoletopoulou et al. find that BDNF-mediated suppression of autophagic flux in neurons is important to promote synaptic plasticity (42). Taken together, BDNF has varying effects on autophagy in neurons, and its precise role in regulating autophagy in axons remain unclear.

Here, we set out to investigate the regulation of axonal autophagy by the neurotrophin BDNF, and elucidate whether autophagosomes carry neurotrophin-mediated signaling information. We find that BDNF induces autophagic flux in primary neurons by stimulating the retrograde pathway for autophagy in axons. We observed an increase in autophagosome density and retrograde flux that culminates in increased autophagosome density in the soma. Surprisingly, we find little evidence of autophagosomes co-migrating with BDNF conjugated to quantum dots (Qdots). BDNF-Qdots robustly co-migrate with retrograde TrkB, but not with LC3-positive autophagic compartments in the axon. Together, we find that BDNF stimulates autophagy in the axon, but retrograde autophagosomes do not appear to carry neurotrophin-based signaling information. Thus, BDNF may exert effects on autophagy as a consequence of BDNF-induced axonal outgrowth and remodeling that necessitates canonical roles for autophagy in degradation, rather than unconventional roles in neurotrophin signaling.

## RESULTS

### BDNF upregulates autophagic flux in primary neurons

Prior studies report varying effects of the neurotrophin BDNF on neuronal autophagy (42,43). Thus, the precise mechanisms by which BDNF affects neuronal autophagy remain unclear. In this study, we define the effects of BDNF on the dynamics of autophagy in neurons in real-time using live-cell imaging, in a compartment-specific manner. First, we assessed the effects of BDNF on total cellular levels of autophagy. For this experiment, we treated primary hippocampal neurons with 50 ng/ml BDNF overnight, and measured levels of autophagy by immunoblotting for endogenous LC3. LC3 has two key forms; a cytosolic form (designated LC3-I) that when lipidated becomes associated with autophagosome membranes (designated LC3-II) (47). These two forms can be distinguished by molecular weight using immunoblot analysis and the amount of LC3-II can serve as a measure of autophagy levels. Since autophagosome production is countered by autophagosome clearance via lysosomes, we also determined the degree of flux through the autophagy pathway by blocking lysosome function using Bafilomycin A1 (Baf A1), an inhibitor of the lysosomal proton pump (48). Baf A1 neutralizes the lysosome, thereby blocking proteolytic activity and autophagosome clearance. In this way, we can more completely capture the total amount of autophagosomes in the cell, and any alterations in response to a given treatment. We found that overnight treatment with BDNF increased levels of LC3-II in neurons by ∼1.9-fold relative to the solvent control (Fig. 1A, B), albeit this effect was not statistically significant. As expected, Baf A1 significantly increased LC3-II levels ∼2.2-fold relative to the solvent control (Fig. 1A, B), indicating an accumulation of autophagosomes due to effective inhibition of lysosome function. Co-treatment of BDNF with Baf A1 further increased LC3-II levels ∼1.7-fold beyond what is accumulated due to Baf A1 alone (Fig. 1A, B), indicating that BDNF stimulates autophagic flux in neurons. Moreover, this increase in LC3-II levels with co-treatment of BDNF with Baf A1 relative to Baf A1 alone reveals that a population of autophagosomes that form in response to BDNF are destined for a lysosomal compartment. Combined, these results demonstrate that neuronal autophagy is stimulated by the neurotrophin BDNF.

**Figure 1.**
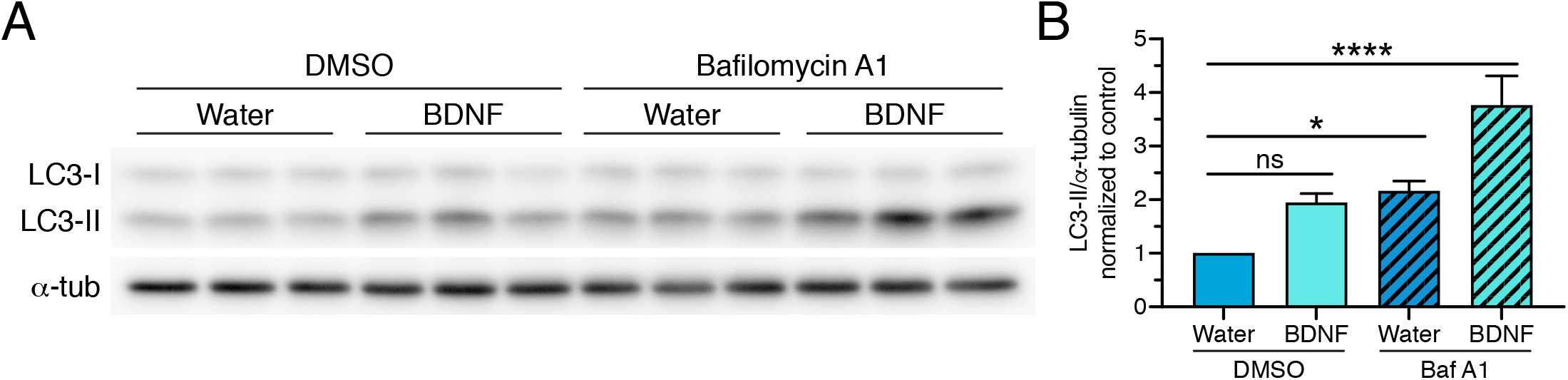
BDNF upregulates autophagic flux in primary neurons. **(A)** Immunoblot analysis and **(B)** corresponding quantification of primary wild type hippocampal neurons treated with 50 ng/ml BDNF (or equivalent volume of water as a solvent control) for 24 hr; 100 nM Baf A1 (or equivalent volume of DMSO as a solvent control) was included in the last 4 hr. LC3-II levels were normalized to α-tubulin (means ± SEM; one-way ANOVA with Dunnett’s post hoc test; n=5 independent experiments; samples in individual experiments were performed in duplicate or triplicate; 9-10 DIV).

Since we previously established that autophagy in neurons is a highly compartmentalized process (24,25,28,49), we next defined the effect of BDNF on neuronal autophagy in a compartment-specific manner using live-cell imaging. We first focused on measuring autophagy levels in the soma since the soma is the final destination for most autophagosomes in the neuron, and houses the majority of lysosome-mediated degradative activity and autophagosome clearance in the neuron (25,50). Thus, by measuring autophagy levels in the soma, we will most effectively capture changes in autophagic flux in response to BDNF. For this experiment, we isolated cortical neurons from transgenic mice expressing GFP-LC3 (51). In this model, autophagosomes are represented by GFP-LC3-positive puncta (Fig. 2A) that we have previously validated to be bona fide autophagic compartments (24,25,28). Neurons were treated overnight with 50 ng/ml BDNF; Baf A1 (or an equivalent volume of DMSO as a solvent control) was included in the final 4 hr to measure autophagic flux. To measure autophagy levels, we performed live-cell imaging using a Spinning Disk confocal microscope and generated Z-stacks spanning the entire depth of the soma. We then generated maximum projections and quantified the area occupied by GFP-LC3-positive puncta using Ilastik, a machine learning-based program to identify and segment objects of interest. We found that BDNF increased the total area of GFP-LC3-positive autophagosomes ∼1.8-fold relative to the solvent control (Fig. 2A, B), albeit this effect was not statistically significant. As expected, blocking lysosome function with Baf A1 increased the total area of GFP-LC3-positive autophagosomes ∼2.8-fold relative to the solvent control (Fig. 2A, B). Co-treatment of BDNF with Baf A1 significantly increased autophagosome area ∼2-fold beyond what is accumulated due to Baf A1 alone (Fig. 2A, B), indicating that BDNF upregulates autophagic flux in neurons.Furthermore, treatment with BDNF, either alone or in combination with Baf A1 resulted in a shift toward greater values for autophagosome area relative to respective controls (Fig. 2C), consistent with increased levels of autophagosomes in response to BDNF. Moreover, our observations that treatment with Baf A1 leads to a larger increase in autophagosome area in BDNF-treated cells compared to water-treated cells indicates that a significant population of autophagosomes that form in response to BDNF are destined for a degradative compartment. Our results with live cell imaging corroborate those from immunoblotting, and indicate that BDNF stimulates autophagic flux in neurons, leading to an accumulation of autophagosomes in the soma.

**Figure 2.**
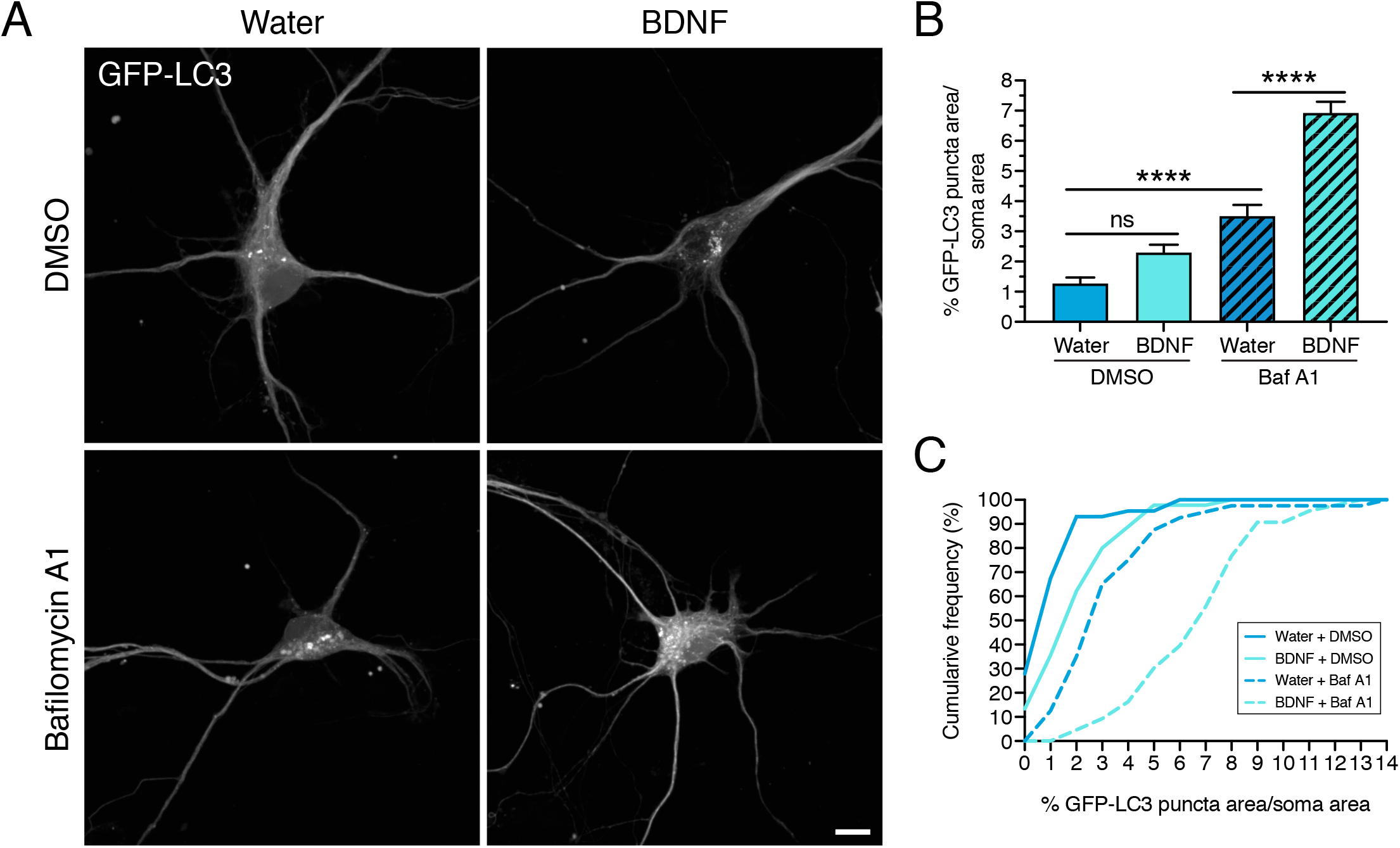
BDNF increases autophagic flux in the soma of primary neurons. **(A-C)** Live-cell imaging of GFP-LC3 transgenic cortical neurons treated with 50 ng/ml BDNF (or equivalent volume of water as a solvent control) for 20 hr; 100 nM Baf A1 (or equivalent volume of DMSO as a solvent control) was included in the last 4 hr. **(A)** Maximum projections of z-stacks of GFP-LC3. Bar, 10 μm. **(B and C)** Corresponding quantitation of the percent area occupied by GFP-LC3-positive puncta normalized to soma area (B, means ± SEM; C, cumulative frequency; one-way ANOVA with Tukey’s post hoc test; n=40-45 somas from 3 independent experiments; 8 DIV).

### BDNF stimulates retrograde transport of axonal autophagosomes

A significant population of autophagosomes present in the soma originate from the axon (24,25,28). We, along with others, found that axonal autophagosomes are produced in axon terminals and delivered to the soma via retrograde transport driven by dynein (23-25,27-31,34-36,52). Since we observed a significant increase in autophagosomes in the soma, we wanted to determine whether BDNF stimulates the retrograde pathway for autophagy in the axon, leading to enhanced delivery of axonal autophagosomes to the soma. To measure the effect of BDNF on axonal autophagy, we treated GFP-LC3 transgenic hippocampal neurons with 50 ng/ml BDNF for overnight, and performed live-cell imaging of axons to track the dynamics of autophagosome motility in real-time. Treatment with BDNF significantly increased the density of autophagosomes in the axon, as measured by the number of GFP-LC3-positive puncta per 100 μm distance along the axon (Fig. 3A, B). Treatment with BDNF also significantly increased the flux of autophagosomes moving in the axon, as measured by the number of autophagosomes that crossed a defined mid-point in the axon per minute (Fig. 3A, C). Lastly, treatment with BDNF significantly increased the percentage of autophagosomes that underwent robust retrograde motility, as defined by the percentage of autophagosomes that traveled a net distance of ≥5 μm during the 5 minute imaging window (Fig. 3A, D). Combined, these results are consistent with a stimulation in autophagosome production in axons and retrograde delivery to the soma in response to BDNF treatment. Thus, we find that BDNF stimulates autophagy in the axons of primary neurons.

**Figure 3.**
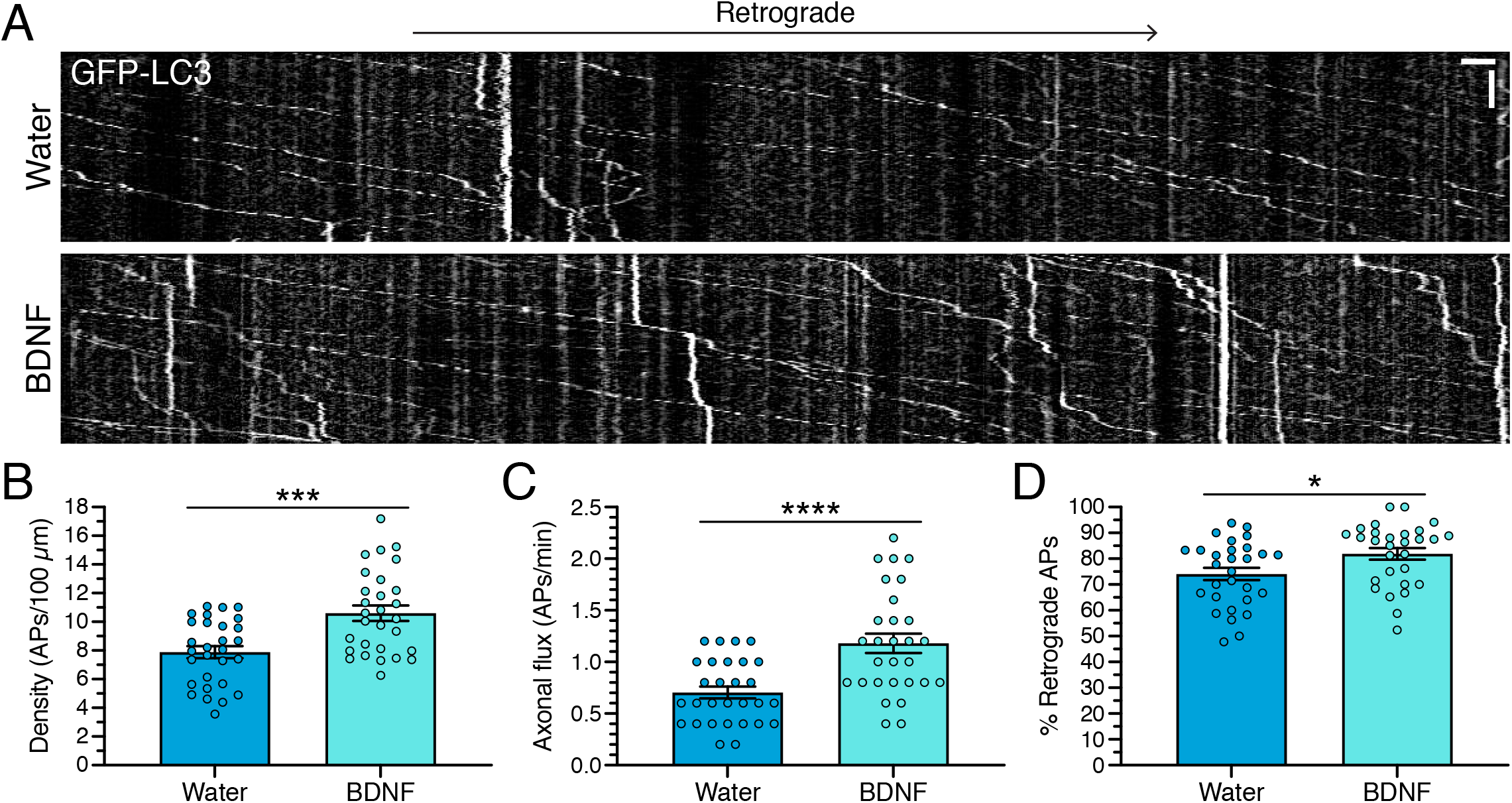
BDNF stimulates retrograde transport of autophagosomes in axons of primary neurons. **(A-D)** Live-cell imaging analysis of GFP-LC3 transgenic hippocampal neurons treated with 50 ng/ml BDNF (or equivalent volume of water as a solvent control) for 24 hr. **(A)** Kymographs of GFP-LC3 motility in the axon; retrograde is to the right. Vertical bar, 1 min. Horizontal bar, 5 μm. **(B)** Quantitation of autophagosome density as measured by the number of autophagosomes per 100 μm of axon (means ± SEM; t-test; n=29 axons from 4 independent experiments; 8-10 DIV). **(C)** Quantitation of autophagosome flux as measured by the number of autophagosomes that cross a defined midpoint in the axon per minute (means ± SEM; t-test; n=29 axons from 4 independent experiments; 8-10 DIV). **(D)** Quantitation of the percentage of autophagosomes that undergo retrograde transport (means ± SEM; t-test; n=29 axons from 4 independent experiments; 8-10 DIV).

### The vast majority of axonal autophagosomes do not transport BDNF cargo

Since we find that BDNF stimulates axonal autophagy, we wondered whether these autophagosomes conveyed signaling information across the axon by transporting BDNF from the distal axon to the soma. To examine this possibility, we determined the degree of overlap between axonal autophagosomes and BDNF conjugated to fluorescent Quantum Dots (605 nm). For this experiment, we incubated neurons with BDNF conjugated to Qdots-605 via a streptavidin-biotin linkage. This method has been widely used to track signaling endosomes in axons (39,53). First, we performed several controls to validate that BDNF was effectively conjugated to the Qdots-605. We transiently expressed TrkB-GFP in wild type neurons and assessed overlap with BDNF-Qdots in the axon using dual-color live cell imaging. As expected, after only a short-term incubation with BDNF-Qdots, we observed robust overlap and co-migration of BDNF-Qdots with TrkB-GFP (Fig. 4A). Strikingly, only the retrograde, and not anterograde, tracks for TrkB-GFP were co-positive for BDNF-Qdots (Fig. 4A), consistent with a selective labeling of retrograde signaling endosomes with BDNF-Qdots. In fact, ∼65% of the retrograde TrkB-GFP-positive organelles were co-positive for BDNF-Qdots (Fig. 4B). In sum, these results indicate that BDNF was effectively conjugated to Qdots-605, and able to successfully bind its receptor, internalize, and undergo retrograde transport in the axon. To further verify specificity of the tracks labeled with BDNF-Qdots, we also incubated neurons with Qdots-605 that were not conjugated to BDNF, but rather incubated with water as a solvent control (Fig. 4A). In contrast to BDNF-Qdots, the Qdots-alone, not conjugated to BDNF, did not exhibit retrograde motility, and did not colocalize with TrkB-GFP (Fig. 4A, B). Thus, the retrograde motility exhibited by BDNF-Qdots is due to specific labeling of organelles transporting BDNF cargo.

**Figure 4.**
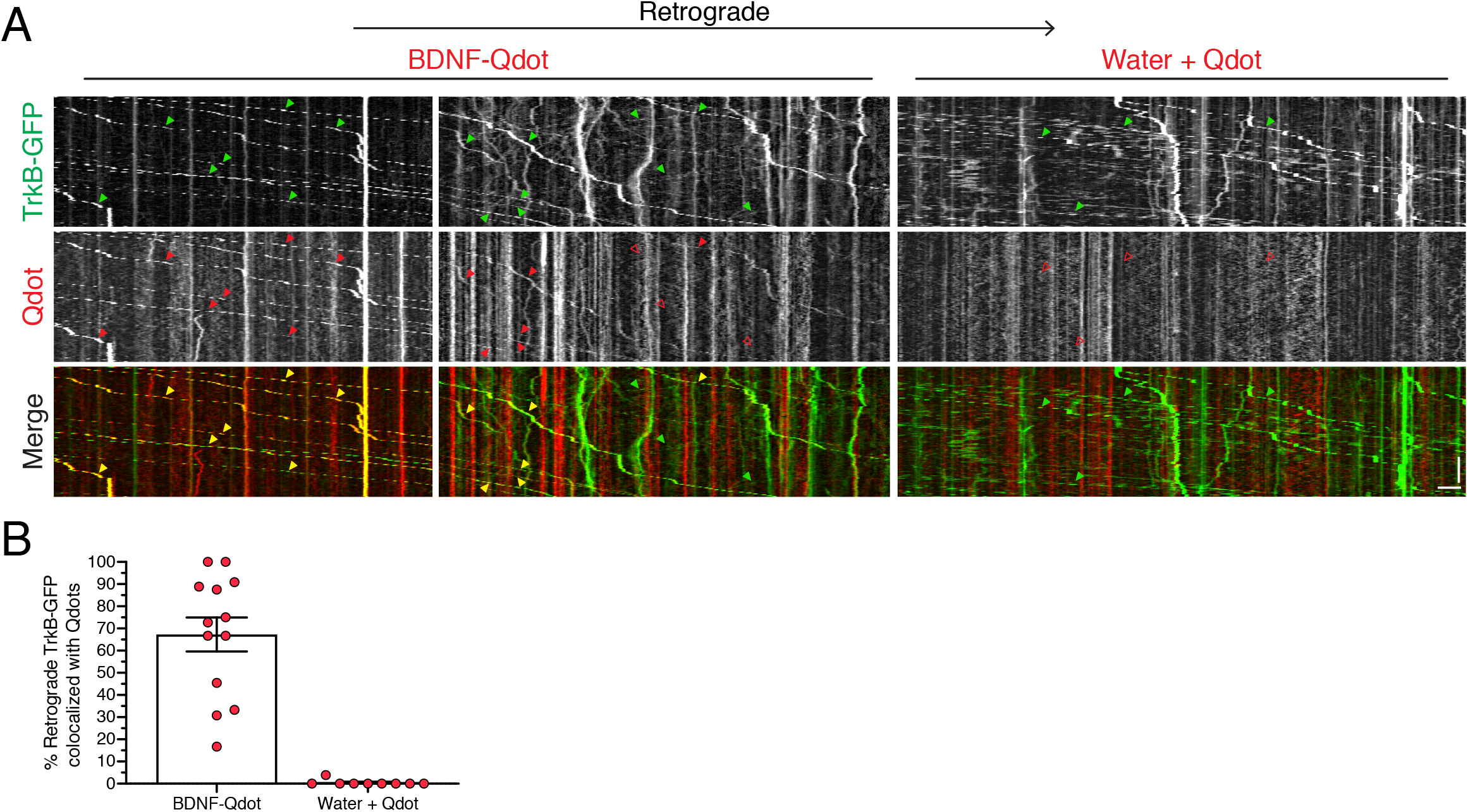
BDNF-Qdots label TrkB-positive signaling endosomes. **(A)** Kymographs of TrkB-EGFP and Qdot (± conjugation to BDNF) motility in the axons of primary wild type cortical neurons after a short-term incubation with Qdots. Throughout the figure, retrograde is to the right. Closed arrowheads denote organelles positive for the respective marker. Open arrowheads denote organelles negative for the respective marker. Closed yellow arrowheads in the merged image denote organelles co-positive for both markers. Vertical bar, 1 min. Horizontal bar, 5 μm. **(B)** Quantitation of the percentage of retrograde TrkB-EGFP-positive organelles that are co-positive for Qdots with or without conjugation to BDNF (means ± SEM; n=9-13 axons from 2 independent experiments; 8-9 DIV).

Next, we wanted to determine whether retrograde autophagosomes in the axon transported BDNF cargo. Thus, we performed dual-color live cell imaging of BDNF-Qdots and GFP-LC3 in axons of primary cortical neurons. First, we incubated neurons with BDNF-Qdots for a short-term treatment lasting ∼3 hrs. Consistent with our prior work, we observed predominantly retrograde transport of autophagosomes in the axon (24,25,28). However, we found very little evidence of colocalization between retrograde autophagosomes and BDNF-Qdots (Fig. 5A, C). In fact, of the 114 autophagosomes counted, only one autophagosome exhibited colocalization with BDNF-Qdot signal (Fig. 5A, C; example of colocalization denoted with yellow arrowhead in the merged kymograph). We did observe retrograde tracks positive for BDNF-Qdots, however, they were negative for GFP-LC3 (Fig. 5A) and likely represented BDNF bound to endogenous TrkB in signaling endosomes. We also examined whether an overnight incubation with BDNF-Qdots might reveal a population of autophagosomes co-positive for BDNF. Similar to the short-term incubation, overnight incubation did not reveal significant overlap between autophagosomes and BDNF-Qdots (Fig. 5B, C). We observed retrograde tracks for GFP-LC3 and BDNF-Qdots, however, these tracks largely did not overlap (Fig. 5B, C). Of the 192 autophagosomes counted, we observed only one event of co-migration between an autophagosome and a BDNF-Qdot. However, further analysis of the movie revealed that these markers were distinct structures that appeared tethered, but were not the same organelle (Fig. 5B; example of tethered LC3 and BDNF-Qdot denoted with closely apposed red and green arrowheads in the merged kymograph). Combined, our data suggest that autophagosomes and compartments positive for BDNF are largely distinct organelle populations in the axon. Furthermore, the vast majority of autophagosomes are negative for BDNF-Qdots indicating that autophagosomes likely do not play a major role in relaying neurotrophic signaling information across the axon in the form of BDNF cargo.

**Figure 5.**
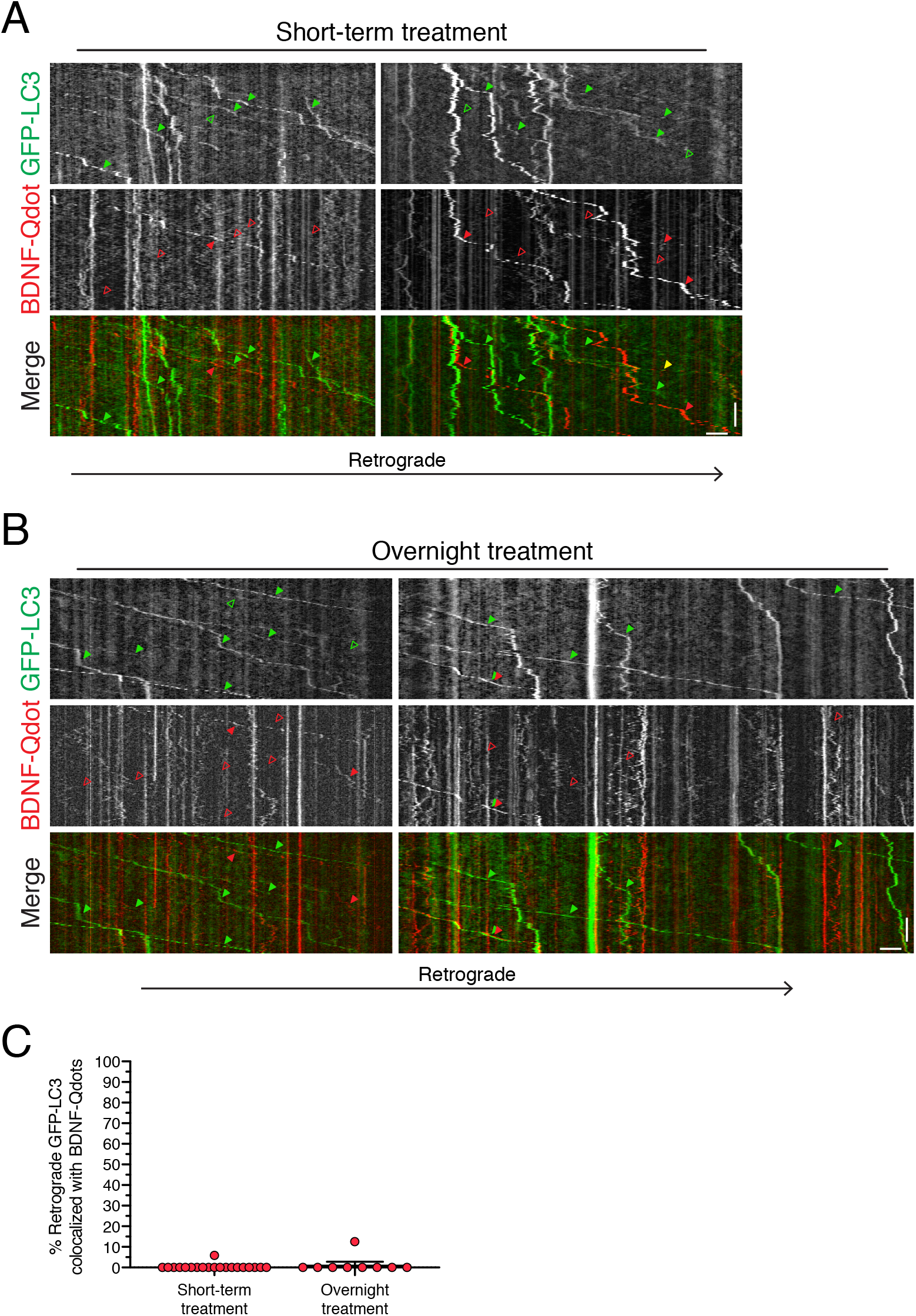
Axonal autophagosomes are not positive for BDNF-Qdots. **(A-B)** Kymographs of GFP-LC3 and BDNF-Qdot motility in the axons of primary cortical neurons after a **(A)** short-term or **(B)** overnight treatment with BDNF-Qdots. Throughout the figure, retrograde is to the right. Closed arrowheads denote organelles positive for the respective marker. Open arrowheads denote organelles negative for the respective marker. Closed yellow arrowheads in the merged image denote organelles co-positive for both markers. Vertical bars, 1 min. Horizontal bars, 5 μm. **(C)** Quantitation of the percentage of retrograde GFP-LC3-positive autophagosomes that are co-positive for BDNF-Qdots (means ± SEM; short-term treatment, n=20 axons from 4 independent experiments, 8 DIV; overnight-treatment, n=9 axons from 2 independent experiments, 8-9 DIV).

## DISCUSSION

Neurotrophins play key roles in regulating the development, function, and survival of axons. But how neurotrophins impact neuronal autophagy, a key pathway that maintains axonal homeostasis, is unclear. We find that BDNF upregulates autophagic flux in primary hippocampal and cortical neurons. BDNF increases global levels of autophagy as measured by immunoblot analysis (Fig. 1), as well as compartment-specific levels in the soma and axon (Fig. 2 and 3). Specifically, BDNF increases the density of autophagosomes in the axon, as well as the number of autophagosomes moving in the retrograde direction (Fig. 3). This stimulation in axonal autophagy delivers more autophagosomes to the soma (Fig. 2). These results, combined with prior observations that a population of axonal autophagosomes are co-positive for TrkB (43,44), led us to examine whether autophagosomes carry signaling information in the form of BDNF. Using dual color imaging of GFP-LC3-positive autophagosomes and BDNF conjugated to Quantum Dots (BDNF-Qdots) we found little evidence of co-migration between these two markers (Fig. 5). By contrast, TrkB-positive organelles were robustly co-positive for BDNF-Qdots (Fig. 4). Moreover, BDNF-Qdots co-migrate with only the retrograde population, and not the anterograde population, of TrkB (Fig. 4). Thus, BDNF conjugated to Qdots is able to effectively bind its cognate TrkB receptor and specifically label signaling endosomes. Lastly, we find that a significant population of autophagosomes that form in response to BDNF are in the process of being targeted to lysosomes for degradation (Fig. 1 and 2). In sum, while BDNF increases autophagy in the axonal compartment of primary neurons, axonal autophagosomes do not appear to carry BDNF-mediated neurotrophin signaling information.

A model proposed in the field is that axonal autophagy may mediate BDNF/TrkB signaling to regulate neuronal development and function (43,44). Kononenko et al. find that the clathrin adaptor complex AP-2, independent of its functions in endocytosis, mediates retrograde transport of autophagosomes positive for TrkB to the soma. Loss of AP-2 reduces neuronal complexity, a phenotype rescued by exogenous BDNF, suggesting that this pathway is important for proper neuronal development (43). Andres-Alonso et al. find that TrkB-positive amphisomes pause at presynaptic boutons to enable local TrkB signaling that influences transmitter release and synaptic plasticity (44). In support of these models, we find that BDNF stimulates the retrograde pathway for axonal autophagy. Our results corroborate those from Kononenko et al. that find that BDNF increases the speed and run lengths of axonal autophagosomes traveling in the retrograde direction (43). However, our observed lack of BDNF in retrograde autophagosomes does not support a major role for autophagy in transporting BDNF/TrkB signals in the axon. Moreover, we, along with several groups, find that nearly all LC3-positive organelles in the axon are positive for LAMP1 (marker for late endosomes and lysosomes), Rab7, and can be tracked with the acidotropic dye, Lysotracker (24,29,31,33,54), suggesting that nearly all autophagosomes in the axon have matured into more acidic amphisomes (fusion products of autophagosomes with late endosomes). Axonal LC3-positive organelles are also co-positive for probes reported to fluoresce with Cathepsin B/D cleavage, raising the possibility that these amphisomes possess some degree of proteolytic activity (32,33). These findings raise questions as to how signaling information might be preserved in the acidic and maturing environment of the amphisome.

Surprisingly, the exact nature of the signaling endosomes that transport BDNF/TrkB continues to be elusive (40,55,56). Evidence shows that TrkB carriers require Rab7 for long-range retrograde transport in axons (57). TrkB carriers also appear to have neutral lumens (58,59) as acidification causes ligand-receptor dissociation (60) and arrests signaling (61). In fact, premature acidification of endosomes attenuates TrkB signaling (62). Mutations in the endosomal Na^+^/H^+^ exchanger 6 (NHE6) that pumps Na^+^ and/or K^+^ into endosomes and H^+^ out of endosomes cause the neurological disorder Christianson syndrome (62). Loss of NHE6 function reduces the proton leak out of endosomes resulting in over-acidifcation of endosomes, precocious activation of cathepsin D in endosomes, increased TrkB degradation, and reduced TrkB signaling (62,63). Consequently, loss of NHE6 reduces branching of dendrites and axons (62). This phenotype can be rescued with exogenous addition of BDNF, suggesting that defects in branching are likely due to attenuated BDNF/TrkB signaling (62). Moreover, modulators of signaling endosome trafficking do not affect the motility of acidic organelles labeled with LysoTracker (e.g. late endosomes and amphisomes), suggesting that the trafficking of signaling endosomes in largely uncoupled from the autophagy pathway in axons (64). Lastly, to potentiate BDNF/TrkB signaling, signaling endosomes must evade lysosomal fate. Suo et al. found that NGF-TrkA signaling endosomes contain the effector protein Coronin-1 which routes signaling endosomes towards a recycling route upon arrival in the soma and prevents fusion with lysosomes (59). Thus, retrograde autophagosomes and signaling endosomes may travel in parallel pathways in the axon, but maintain distinct identities.

We propose that BDNF stimulates neuronal autophagy for its canonical functions in degradation. BDNF functions as a short-range signal that regulates axon terminal arborization, and synaptic development and maintenance at the target (65). Thus, BDNF-induced processes involve the turnover of proteins, membranes, and organelles that may require an induction of autophagy. In fact, loss of core autophagy genes (e.g. *Atg7* and *Atg5*) can result in an accumulation of smooth ER in presynaptic terminals, suggesting an important role for autophagy in membrane turnover and recycling in distal axons. (14,16). More recent studies have shown that autophagy negatively regulates presynaptic neurotransmission by controlling the size of the ER in presynaptic terminals (7). Autophagy can also modulate presynaptic neurotransmission by reducing synaptic vesicle pool size and release probability (8,9). BDNF is also an important regulator of synaptic plasticity and can act at presynaptic sites (66), but whether BDNF regulates synaptic development, function, and plasticity through autophagy, and how this would be achieved, is only beginning to be understood. In specific contexts, BDNF may suppress autophagy *in vivo* to promote synaptic plasticity (42). Moreover, the in vitro nature of our studies cannot address how neighboring glia may impact these processes. Future studies will need to unravel the connections between BDNF/TrkB and their impacts on the various functional modalities of autophagy.

### EXPERIMENTAL PROCEDURES

#### Reagents

Transgenic mice expressing GFP-LC3 (EGFP fused to the C-terminus of rat LC3B) were obtained from the RIKEN BioResource Research Center (RBRC00806; strain name B6.Cg-Tg(CAG-EGFP/LC3)53Nmi/NmiRbrc; GFP-LC3#53) ([Mizushima et al, 2004](https://pubmed.ncbi.nlm.nih.gov/14699058/)) and maintained as heterozygotes. All animal protocols were approved by the Institutional Animal Care and Use Committee at the University of Pennsylvania. Primary antibodies for immunoblot analysis include rabbit anti-LC3 (Abcam, Ab48394) and mouse anti-TUBA1A/α-tubulin (Sigma, T-9026). Peroxidase conjugated secondary antibodies include donkey anti-rabbit (Jackson ImmunoResearch Laboratories, 711-035-152), and donkey anti-mouse (Jackson ImmunoResearch Laboratories, 715-035-151). Reagents include Bafilomycin A_1_ (Sigma, B1793), Human BDNF (Alomone Labs, B-250), Human BDNF-Biotin (Alomone Labs, B-250-B), and streptavidin-conjugated Quantum Dots 605 (ThermoFisher Scientific, Q10103MP). TrkB-mEGFP (Addgene plasmid # 83952) was expressed in primary neurons using Lipofectamine 2000, following the manufacturer’s protocol.

#### Primary Neuron Culture

Wild type mouse hippocampal neurons for immunoblot analysis were obtained from the Neurons R Us core facility at the University of Pennsylvania and prepared from C57BL/6 mouse embryos at gestational day 18. For live-cell imaging experiments, GFP-LC3 transgenic mouse hippocampal and cortical neurons were isolated from GFP-LC3–transgenic mouse embryos at gestational day 15 as previously described; Fig. 4 uses non-transgenic neurons isolated from non-transgenic littermates ([Dong et al., 2019](https://www.ncbi.nlm.nih.gov/pmc/articles/PMC8020715/#bib5); [Kulkarni et al., 2020](https://www.ncbi.nlm.nih.gov/pmc/articles/PMC8020715/#bib26)). In brief, hippocampal or cortical tissue was digested with 0.25% trypsin for 10 min at 37°C, and then triturated to achieve a homogeneous cell suspension. For imaging experiments, neurons were plated at a density of either 750,000 or 1,500,000 neurons per 10-cm dish (1.2 or 2.4 × 10^4^ neurons/cm^2^ respectively) filled with eight 25-mm acid-washed glass coverslips coated with 1 mg/ml poly-L-lysine (PLL). To achieve single neuron resolution, GFP-LC3 transgenic neurons were diluted 1:20 with non-transgenic neurons. For immunoblotting experiments, 240,000 (∼2.4 × 10^4^ neurons/cm^2^) neurons were plated per well of a 6-well plate coated with 0.5 mg/ml PLL. Neurons were grown for 8-10 DIV in mouse neuron maintenance medium (neurobasal medium supplemented with 2% B-27, 33 mM glucose, 37.5mM NaCl, 2mM GlutaMAX, 100 U/ml penicillin, and 100 μg/ml streptomycin) at 37°C in a 5% CO_2_ incubator. Every 3–4 d, 20–30% media was replaced; 1 μM AraC (anti-mitotic drug; Sigma, C6645) was added to the first feed.

#### BDNF Treatment

For BDNF treatments, neurons were incubated in mouse maintenance media supplemented with 50 ng/ml of Human BDNF, or an equivalent volume of H_2_O as a solvent control, for 18-24 hours. In Fig. 1 and 2, media was supplemented with 100 nM Bafilomycin A_1_ or an equivalent volume of DMSO as a solvent control during the last 4 hours of BDNF or H_2_O treatment.

#### Immunoblotting

Following BDNF treatment, neurons were washed in PBS (150 mM NaCl, 50 mM NaPO_4_, pH 7.4) and lysed in RIPA buffer (150 mM NaCl, 1% Triton X-100, 0.5% deoxycholate [Thermo Fisher Scientific, BP349-100], 0.1% SDS [Thermo Fisher Scientific/Invitrogen, 15-553-027], 1X complete protease inhibitor mixture supplemented with EDTA [Sigma, 11697498001], 50 mM Tris-HCl, pH 7.4) for 30 min on ice. Following lysis, samples were centrifuged at 17,000 x g for 15 min at 4°C. Supernatants were analyzed by SDS-PAGE and transferred onto an Immobilon-P PVDF membrane. Membranes were blocked in 5% milk in TBS-Igepal (2.7 mM KCl, 137 mM NaCl, 0.05% Igepal [Sigma, I3021], 24.8 mM Tris-HCl, pH 7.4) for 30 min at room temperature, followed by incubation in primary antibody diluted in block solution for overnight at 4°C, rocking. The following day, membranes were washed 3 times for 20 min each in HRP wash buffer (150 mM NaCl, 0.1% BSA, 0.05% Igepal, 50 mM Tris-HCl, pH 8.0) and incubated in peroxidase-conjugated secondary antibody diluted in HRP wash buffer for 45 min at room temperature, rocking. Following incubation in secondary antibody, membranes were washed 3 times for 20 min each in HRP wash buffer and developed using the SuperSignal West Pico Chemiluminescent Substrate (Thermo Fisher Scientific, PI34580).

#### Live cell Imaging

##### Microscope

Coverslips were sandwiched within a Chamlide CMB magnetic imaging chamber (BioVision Technologies). Live-cell imaging was performed on a BioVision spinning-disk confocal microscope system with a Leica DMi8 inverted widefield microscope, a Yokagawa W1 spinning-disk Confocal microscope, and a Photometrics Prime 95B scientific complementary metal–oxide–semiconductor camera. Images were acquired with the VisiView software using a 63×/1.4 NA plan apochromat oil-immersion objective and solid-state 488nm and 561nm lasers for excitation of GFP-tagged proteins and Qdots-605, respectively. For images that would be quantitatively compared directly to each other, the same acquisition parameters were used across treatment conditions. The microscope is equipped with an adaptive focus control to maintain a constant focal plane during live-cell imaging, and an environmental chamber to maintain temperatures at 37°C.

##### Autophagic flux

For live-cell imaging of GFP-LC3 (Fig. 2 and 3), samples were washed once in Hibernate E supplemented with 2% B-27 and 2 mM GlutaMAX (HibE imaging solution) and imaged in the same solution. In Fig. 2, neurons were imaged in HibE imaging solution supplemented with either BDNF or H_2_O, and 100 nM Bafilomycin A_1_ or equivalent volume of DMSO as a solvent control. Z-stacks spanning the entire depth of the soma were acquired at 0.2 μm sections. In Fig. 3, neurons were imaged in HibE imaging solution and single GFP-LC3 transgenic axons were identified and imaged in the mid-axon, as defined by >100 μm from the distal end of the axon and soma. Images were acquired at one frame every 2 seconds for 5 minutes. Each coverslip was imaged for a maximum of 40 minutes on the spinning disk confocal microscope in an environmental chamber at 37°C.

##### BDNF-Qdots

BDNF-biotin (54 nM) and streptavidin-conjugated Qdots-605 (40 nM) were conjugated in neurobasal media for 1 hour on ice. To deplete neurons of endogenous BDNF, neurons were washed 2-3 times for 5 minutes each wash with fresh neurobasal medium (Gibco, 21103-049). BDNF-Qdots were then diluted 1:200 in fresh neurobasal media and then added to neurons. In Fig. 4, primary mouse cortical neurons were transfected with TrkB-mEGFP using Lipofectamine 2000 two days prior to incubation with BDNF-Qdots. At 8-9 DIV, neurons were incubated as described above with BDNF-Qdots or with a Qdot-only control (equal volume of H_2_O was substituted for BDNF-biotin during the conjugation for 1 hr on ice) for 3-7 hours. In Fig. 5, primary mouse cortical neurons were incubated with BDNF-Qdots for either 2.5-3.5 hours (short-term) or 16-23 hours (long-term). To prevent detrimental effects of sustained nutrient deprivation in the long-term treatment, neurons were washed in mouse maintenance media, and incubated with BDNF-Qdots diluted in mouse maintenance media. For imaging short-term and long-term treatments, neurons were washed and imaged in HibE imaging media; images were acquired at 1 frame every 2 seconds for 5 minutes. Samples were imaged for a maximum of 50 minutes on the microscope.

#### Image Analysis

##### Immunoblotting Analysis

Bands were quantified using the gel analyzer tool in Fiji. Boxes were drawn around each individual band and plotted on a densitometry graph. After plotting the densitometry graphs, areas under the curve were measured using the magic wand tool in Fiji. To control for sample loading differences between lanes, values for LC3-II were divided by values for TUBA1A/α-tubulin from each corresponding lane. Values were then normalized relative to the control sample (mouse maintenance media with H_2_O and DMSO) and expressed as a fold difference above the control.

##### Analysis of Autophagic Flux in the Soma

Maximum projections of Z-stacks were generated in Fiji, and the neuron soma was outlined and measured for total cross-sectional area. Ilastik, a machine learning-based program, was trained to identify and segment GFP-LC3-positive puncta based on the intensity, edge gradients, and texture of the signal. Ilastik segmentations were imported into Fiji and the areas of identified objects were measured using the “analyze particles” function. Total area occupied by GFP-LC3-positive puncta in the soma was normalized to the respective soma area, and expressed as a percentage.

##### Analysis of Autophagy in the Axon

In BDNF- or water-treated neurons, GFP-LC3 transgenic axons that had at least one GFP-LC3-positive punctum that traveled a net distance of ≥5 μm were selected for analysis. Kymographs from the mid-axon of selected axons were generated using the MultipleKymograph plugin in FIJI, using a line width of 3 pixels. Using these kymographs and the associated movie, the total number of autophagosomes in the first frame was determined and normalized by kymograph length, and reported as autophagosome density (Fig. 3B). Axonal flux reported in Fig. 3C was determined by the number of vesicles in the kymograph moving past the midpoint of the kymograph, and normalized by duration of movie. Lastly, from each kymograph, the number of vesicles moving in the net retrograde direction (≥5 μm) or the net anterograde direction (≥5 μm) was determined. Non-processive vesicles that did not move a net distance of 5 μm were binned as exhibiting bidirectional and stationary motility. Vesicles from all three bins (retrograde, anterograde and stationary/bidirectional) were summed, from which the percent retrograde vesicles were reported in Fig. 3D.

##### BDNF-Qdot Colocalization

Axons that contained at least one TrkB-EGFP-positive (Fig. 4) or GFP-LC3-positive (Fig. 5) punctum that traveled a net distance of ≥5 μm in the retrograde direction were selected for analysis. Kymographs of selected axons were generated >100 μm from the distal end of the axon using the KymoToolBox Fiji plugin and a line width of 3 pixels. From these kymographs, only retrograde organelles, defined as puncta traveling a net distance of ≥5 μm in the retrograde direction, were included in the analysis. To determine co-localization with BDNF-Qdots, all retrograde TrkB-EGFP (Fig. 4) and GFP-LC3 (Fig. 5) puncta were first counted from each kymograph, and then binned as colocalized if they overlapped and co-migrated with a BDNF-Qdot punctum throughout the duration of the movie. The percentage of GFP-LC3 or TrkB-EGFP puncta that colocalized with BDNF-Qdots was determined for each individual kymograph and based on an average of ∼6-13 vesicles per axon across treatment conditions.

#### Statistical Analysis and Figure Preparation

All image measurements were obtained from the raw data. GraphPad Prism was used to plot graphs and perform statistical analyses; statistical tests are denoted within each figure legend (ns, not significant; *p ≤ 0.05; ***p ≤ 0.001; ****p ≤ 0.0001). For presentation of images, maximum and minimum gray values were adjusted linearly in FIJI and images were assembled in Adobe Illustrator.

## ACKNOWLEDGMENTS

This work was supported by NIH grant R01NS110716 and associated Research Supplement to Promote Diversity in Health-Related Research, NIH grant R00NS082619, the McCabe Fund Fellow Award, the University of Pennsylvania Alzheimer’s Disease Core Center, the Intellectual and Developmental Disabilities Research Center at the Children’s Hospital of Philadelphia and the University of Pennsylvania, and the Philadelphia Foundation to SM. We thank members of the Maday lab for constructive feedback on the manuscript.

